# X-Tracker: Automated Analysis of Xenopus Tadpole Visual Avoidance Behavior

**DOI:** 10.1101/2024.10.10.617688

**Authors:** Caroline R. McKeown, Aaron C. Ta, Christopher L. Marshall, Natalie J. McLain, Kailey J. Archuleta, Hollis T. Cline

## Abstract

*Xenopus laevis* tadpoles exhibit an avoidance behavior when they encounter a moving visual stimulus. A visual avoidance event occurs when a moving object approaches the eye of a free-swimming animal at an approximately 90-degree angle and the animal turns in response to the encounter. Analysis of this behavior requires tracking both the free-swimming animal and the moving visual stimulus both prior to and after the encounter. Previous automated tracking software does not discriminate the moving animal from the moving stimulus, requiring time-consuming manual analysis. Here we present X-Tracker, an automated behavior tracking code that can detect and discriminate moving visual stimuli and free-swimming animals and score encounters and avoidance events. X-Tracker is as accurate as human analysis without the human time commitment. We also present software improvements to our previous visual stimulus presentation and image capture that optimize videos for automated analysis, and hardware improvements that increase the number of animal-stimulus encounters. X-Tracker is a high throughput, unbiased, and significant time-saving analysis system that will greatly facilitate visual avoidance behavior analysis of *Xenopus laevis* tadpoles, and potentially other free-swimming organisms. The tool is available at https://github.com/ClineLab/Tadpole-Behavior-Automation.

## Introduction

Animal behavior is a valuable endpoint to assess brain function in healthy and disease states. From insects to rodents to non-human primates, behavioral outputs have been instrumental in assessing brain function and exploring the underlying neuronal circuitry (Bala et al., 2020; Budick and O’Malley, 2000; Gal et al., 2020; Hubel and Wiesel, 1959, 1962, 1968;Lust and Tanaka, 2019; Neuhauss, 2003; Niell, 2015; Portugues and Engert, 2009; Storchi et al., 2020; Zhu, 2013). Behaviors in frogs have been used extensively to evaluate brain function. For instance, altered fly catching behavior following the famous eye-rotation experiments by Roger Sperry demonstrated fundamental principles of visual system connectivity (Sperry, 1944), and similar studies in the African clawed frog *Xenopus laevis* demonstrated the impact of visual input on visual circuit plasticity (Udin, 1985). More recently, establishment of a range of reproducible and quantifiable behaviors in *Xenopus* tadpoles has greatly expanded the use of this experimental system (Dong et al., 2009; Gambrill et al., 2019; Khakhalin et al., 2014; Liu and Cline, 2016; Liu et al., 2018; McKeown et al., 2013; Pratt and Khakhalin, 2013; Shen et al., 2014; Shen et al., 2011; Truszkowski et al., 2016). In the developing *Xenopus* tadpole, visually-guided behaviors have been particularly useful in demonstrating adverse effects on brain function in models of neurodevelopmental disorders such as autism, seizure, and brain injury (Bell et al., 2011; James et al., 2015; Liu and Cline, 2016; McKeown et al., 2013; Spawn and Aizenman, 2012; Truszkowski et al., 2016). Tadpoles exhibit a visual avoidance behavior when they encounter a moving stimulus (Dong et al., 2009; McKeown et al., 2013; Shen et al., 2011). Alterations to visual system circuitry by genetic manipulation (Gambrill et al., 2019; Liu and Cline, 2016; Liu et al., 2018; Shen et al., 2014; Truszkowski et al., 2016), drug treatment (James et al., 2015; Khakhalin et al., 2014; Spawn and Aizenman, 2012), or brain injury (McKeown et al., 2013) result in deficits in the visual avoidance response, which can be reversible, highlighting the importance of studying these behaviors to neurodevelopment research.

Despite the value of behavioral assays in neuroscience research (Datta et al., 2019; Krakauer et al., 2017; Tully et al., 1994), analysis of the complex datasets collected during naturalistic behavior experiments has been challenging and time-consuming, potentially involving subjective investigator-based analysis (Mathis et al., 2018). Automated analysis of behavior including optimized video and other quantifiable endpoints, as well as computer-based analysis of datasets, increases the efficiency and reproducibility of behavioral assays (Bala et al., 2020; Batty E, 2019; Gal et al., 2020; Storchi et al., 2020). Moreover, high-resolution imaging coupled with automated analysis has resulted in the identification of new behaviors that would otherwise have been missed by the human experimenter (Batty E, 2019; Card and Dickinson, 2008; Creton, 2009; Pereira et al., 2019; Storchi et al., 2020). Nonetheless, the immense power of behavioral research is often bypassed due to the time commitment of the experiments and the complexity of the data (Krakauer et al., 2017), a problem that can be addressed by automated analysis of behavior data (Datta et al., 2019; Gal et al., 2020; Storchi et al., 2020).

Here we present X-Tracker, an automated behavior tracking code that can detect and discriminate moving visual stimuli and free-swimming animals and score encounters and avoidance events. We also present software improvements to our previous visual stimulus presentation and video capture to optimize image data for automated analysis and hardware improvements that increase the number of animal-stimulus encounters. X-Tracker increases resolution of specific events contributing to the behavior, speeds analysis and decreases inter-investigator variability compared to manual analysis. X-Tracker is as accurate as manual human analysis without the human time commitment. These tools greatly facilitate visual avoidance behavior analysis of *Xenopus laevis* tadpoles, and potentially other free-swimming organisms.

## Methods

### Animals

Albino *Xenopus laevis* tadpoles of either sex were obtained by in-house breeding or purchased from Xenopus Express (Brooksville, FL). Tadpoles were reared in vivarium water (pH 7.0), on a 12h:12h light-dark cycle at 22°C. Tadpoles were assessed for visual avoidance behavior at stage 47 (Nieuwkoop and Faber, 1967), as described previously (McKeown et al., 2013). We found that animals are more likely to swim if there are other animals present in the chamber, therefore we test 5 animals per video in open field tests and 2 animals per lane in the channel chamber tests (Table 1). All animal protocols have been approved by the Institutional Animal Use and Care Committee of the Scripps Research Institute.

**Table 1.** Comparison of Parameters between different behavior analysis methods. *(*typical experiment)*

|  | Open Field Manual | Lap Channel Manual | X-Tracker Automated |
| --- | --- | --- | --- |
| Video Length (frames) | 825-975 | 825-975 | 1800 |
| Video Length (seconds) | 55-65 | 55-65 | 60 |
| Animals per video | 5 | 10 | 10 |
| Animals per condition* | 20 | 20 | 20 |
| Videos per condition per day | 4 | 2 | 2 |
| Conditions per experiment* | 5 | 5 | 5 |
| Timepoints per experiment* | 8 | 8 | 8 |
| Total videos per experiment* | 160 | 80 | 80 |
| Total animals per experiment* | 100 | 100 | 100 |
| Events scored per animal | 10 | 10 | unlimited/all |
| Events scored per experiment* | 8,000 | 8,000 | unlimited/all |
| Analysis time per animal | ~70s | ~30s | 2min |
| Analysis time per video | ~6min | ~5min | 20min |
| Analysis time per experiment* | ~16h | ~6.7h | n/a |
| Biological replicates* | 3 | 3 | 3 |
| Total analysis time | ~48h | ~20h | n/a |
| Total hands-on analysis time | ~48h | ~20h | ~30s |

### Behavior Rig Specifications and System Requirements

#### Chamber

Channel chambers were machined out of 8mm thick optically transparent acrylic. The chambers are 108mm × 165mm (4.25 × 6.5 inches) with 5 evenly spaced grooves (spaced 10mm apart) that are 8mm wide and 6mm deep, curving up to 4mm deep on the sides. To avoid diffraction and shadows from the well edges, it is advisable to fill the wells to the rim for behavior assays.

#### Lighting

To visualize tadpoles, the behavior chamber is backlit with 4 infrared (850nm/ 940nm) Single Chip Flexible LED lighting strips (SMD3528-600-IR, LED Lights World, Huake Limited, Guangdong, China) evenly spaced below the platform and flanking the projector (Figure 1A,B).

**Figure 1.**
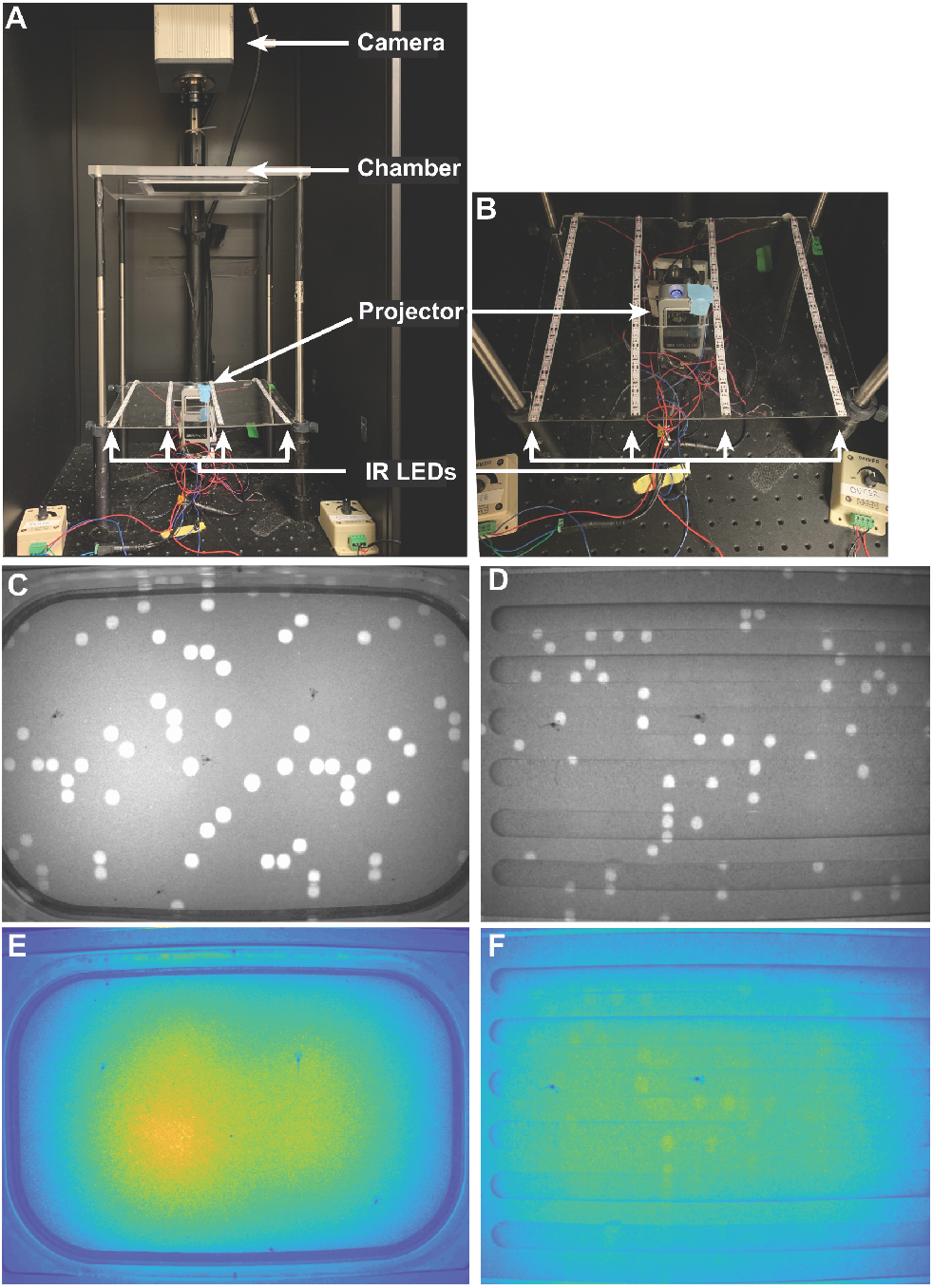
Xenopus visual avoidance behavior apparatus. A) Behavior rig set-up showing the imaging chamber in the middle with the projector and lighting below and the camera above to record video. B) Zoom of A showing the platform with IR LED lighting strips. Note that the lighting has to be placed slightly below the projector so as not to interfere with the visual stimulus projection. C) Image of Open Field assay showing 6 tadpoles in the arena and dot stimuli projected on the bottom of the chamber. This image was taken showing the IR LED arrays used previously (McKeown et al., 2013). D) Image of the Channel assay showing 6 tadpoles, 2 per channel, and dot stimuli projected on the bottom of the chamber. This image was taken using the new IR LED strips. E) Heat map of the lighting from the Open Field image shown in C after filtering out the dots. Note the uneven field illumination from the IR LED arrays. F) Heat map of the lighting from the Channel image shown in D after filtering out the dots. Note the improved evenness of field illumination using the distributed IR LED strips.

#### Projector

Visual stimuli are presented using a microprojector (3M, MPro110) positioned below a clear Plexiglass platform fitted with a translucent sheet of 3M (St. Paul, MN) projector screen (Figure 1A,B), as described previously (Gambrill et al., 2019; McKeown et al., 2013; Shen et al., 2011).

#### Presentation Computer

For presenting the visual stimulus, a Windows 10 or later computer, capable of running Matlab 2014b and Psychtoolbox-3, is required. Matlab 2014b installations of Psychtoolbox-3, Image Acquisition Toolbox, and Image Processing Toolbox software are all required to run the stimulus presentation. The system must be equipped with a NVIDIA graphics card to control the camera through Matlab using

Psychtoolbox-3. At least 12 GB of RAM is recommended to run the stimulus presentation program.

#### Camera

X-Tracker is designed to utilize the Hamamatsu Electron Multiplier CCD Digital Camera C9100-50, which captures 28.1 frames per second at 1000×1000 pixel resolution. The DCAM-API Firebird Phoenix driver allows it to interface with the computer, while the Hamamatsu adaptor for Image Processing Toolbox allows it to be controlled by Matlab. If another camera is utilized, it must be able to record at least 28-30 frames per second and be supported by Image Processing Toolbox; a list of supported cameras and required adaptors, if any, are listed on the Matlab website.

#### Analysis Computer

To run X-Tracker, a Windows 7 or later computer, capable of running Matlab R2017b, is required. The system needs the Matlab Image Processing Toolbox to extract and analyze video frames. At least 32 GB of RAM are recommended, particularly when frequently utilizing the batch input functionality.

#### User Instructions

A detailed user manual is included in the Cline Lab GitHub repository. Briefly, after performing behavior assays, users will compile video data into a single folder. Then users will launch X-Tracker in this folder using MatLab 2017b and select the X-Tracker code. The user will be asked to input selections for tadpole eye-gut distance, avoidance angle tolerance, speed and direction variability, and detection threshold settings, and then run the program. The code will provide progress updates but is completely hands-free from this point. Data will be deposited in the form of Microsoft Excel spreadsheet file (.xls) and saved into the original data folder. For detailed instructions, please see the User Manual in the ReadMe document in the Cline Lab GitHub repository at https://github.com/ClineLab/Tadpole-Behavior-Automation.

### Results

We will first describe the typical experimental design and analysis of visual avoidance behaviors currently used and then describe how we implemented improvements. We use visual avoidance behavior as an assay for visual system function in *Xenopus laevis* tadpoles (Dong et al., 2009; Gambrill et al., 2019; McKeown et al., 2013; McKeown et al., 2017; Shen et al., 2011). In typical behavior experiments, groups of 5 tadpoles were placed in a clear plexiglass open field chamber fitted with a translucent 3M projector screen. Visual stimuli of randomly arrayed 0.4cm diameter moving dots were created by a custom-written MatLab code and presented with a microprojector below the chamber (Figure 1A). Tadpoles were visualized with an array of infrared LEDs and video recordings of tadpole movements and visual stimuli were captured with a Hamamatsu ORCA-ER digital camera (McKeown et al., 2013). The entire system is enclosed in a light-tight compartment.

Tadpole visual avoidance behavior was analyzed manually by post-hoc frame-by-frame viewing and scoring of encounters and avoidance responses. An encounter is defined as a dot moving perpendicularly (within 90±15 degrees) across the tadpole’s eye and an avoidance response is scored when a tadpole displays a sharp turn within 500ms of an encounter (McKeown et al., 2013; Shen et al., 2011). Animals swim in an open field chamber, making the random occurrence of a 90 degree encounter rare. We scored responses to 10 encounters per animal; therefore, animals were filmed for 1 minute to ensure that at least 10 encounters occurred. Because of clutch-to-clutch variation, every experiment included untreated control animals which also allowed evaluation of overall clutch health. Experiments include multiple timepoints per animal and are repeated at least 3 times from independent clutches. Overall, a typical experimental design requires hundreds of animals per experiment, with thousands of encounters and avoidance responses counted (Table 1).

In an effort to simplify and automate this visual avoidance behavior analysis, we have made 3 significant alterations to our behavior assay: 1) modified the behavior chamber to increase the number of encounters, 2) modified the image capture resolution and lighting to increase the quality and consistency of the videos, and 3) developed software to track both animals and dots and automatically score avoidance events. These are described below.

### Behavior chamber modifications – Channel System

Tadpole eyes are on the sides of their heads and they respond most consistently to looming visual stimuli, a dot, approaching the eye perpendicularly (Dong et al., 2009; Khakhalin et al., 2014). Consequently, scoring visual avoidance behavior in free-swimming tadpoles in large open field chambers was inefficient because requisite encounters were relatively rare (Figure 1C). To increase the frequency of tadpole encounters with the dot stimuli, we created a chamber with channels that force animals to swim in a trajectory that is always perpendicular to the moving stimuli. The channel chamber is a custom 108mm × 165mm block of acrylic machined with 5 evenly-space grooves that are 140mm long, 8mm wide, and 6mm deep (Figure 1D). The width of the channels is large enough for a tadpole to execute an avoidance response turn and is sufficient for 2 tadpoles to pass each other (Figure 2B). Because tadpoles tend to remain stationary in sharp corners, the channels were machined with a rounded bottom (8mm curving up to 6mm) and rounded ends (140mm long channels curving to 135mm in length) to encourage the tadpoles to continue swimming. We observed no obvious change in overall swimming behavior between animals in the channel chambers versus animals in the open field chamber, and animals in the channels were confirmed to exhibit an avoidance response to an approaching visual stimulus, detected as a rapid change in trajectory (Figure 2A, B). To evaluate whether the channels alter behavior, we tested the avoidance behavior of the same group of animals in both the open field and channel system over the course of several days, and scored avoidance events using manual analysis. While we saw a significant increase in the number of scorable encounters per animal and the number of scorable animals per video using the channel chamber, we saw no significant difference in the average avoidance responses between the open field and the channels (Figure 2C). By directing the animals to move perpendicularly to the visual stimuli, the channels increase the number of scorable encounters, resulting in both increased speed and ease of analysis (Table 1). These data indicate that the channel chamber does not affect swimming or avoidance behaviors but does increase encounters, simplifying the analysis.

**Figure 2.**
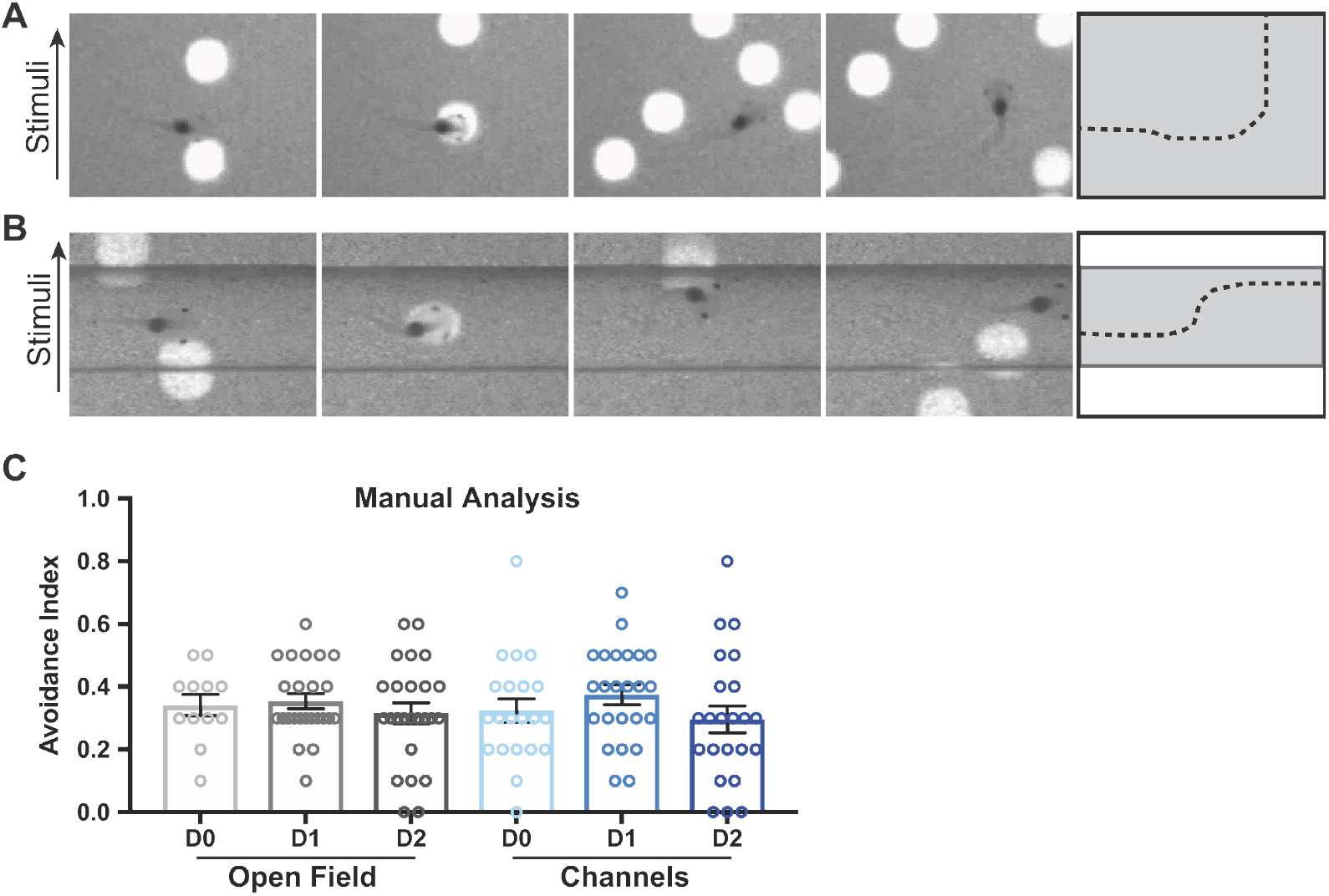
Channel system vs. Open Field for animal behavior. A) Example avoidance response in the Open Field chamber. The animal is swimming toward the right when it visually encounters a moving dot approaching from the bottom of the image. The panels are shown in succession over 500ms total time. A trace of the avoidance response is provided in the last panel. B) Example avoidance response in the Channel system. The animal is swimming toward the right when it visually encounters a moving dot approaching from the bottom of the image. The panels are shown in succession over 500ms total time. A trace of the avoidance response is provided in the last panel. Note that the dots are presented as the same size (0.2um radius) in both systems, however they appear different due to camera differences described in the text, and likely due to differences in the density and refractive index of the acrylic chambers. C) Manual analysis of avoidance indices comparing Open Field (grays) vs. Channels (blues) over several days in the same group of animals. Data are shown as average ± SEM overlaid with individual data points. There is no significant difference between the groups by ANOVA with post hoc Tukey’s for multiple comparisons.

### Visual stimuli presentation/capture modifications

Previously, we used 4 IR LED arrays to backlight the behavior chamber to visualize the tadpoles. This lighting system is sufficient for manual behavior analysis, but it creates an uneven field illumination as shown by the heat map generated in Matlab (Figure 1E). This uneven illumination is problematic for automated tracking software that relies on fixed thresholding to identify objects. We created a more uniform illumination using 4 evenly spaced

Infrared LED strips (SMD3528-600-IR, LED Lights World) (Figure 1B, F). In addition, we improved the captured images by upgrading the camera to a Hamamatsu C9100-50 which has increased sensitivity and resolution, and a higher frame rate of 28.1 frames per second. Utilizing the MP4 instead of AVI video container format enabled us to improve the camera capture resolution of recorded videos from 80% to 95% quality while maintaining a manageable file size of 120MB.

Changing out the camera required a concurrent change to the computer and the stimulus presentation and capture code. The computer hardware specifications are listed in the System Requirements section below and the Stimulus Presentation Code is available on the Cline Lab GitHub. Lastly, we modified the acquisition settings to collect a fixed number of 2250 frames of video in order to standardize the video length for analysis; the initial 450 frames are not scored by X-Tracker to ensure consistency regarding stimulus presentation start-up time (Table 1). Together these modifications allow for a higher quality uniform image which creates an optimal input for automated detection software.

### X-Tracker: Automated Behavior Analysis Software

To automate the visual avoidance behavior analysis, we were tasked with creating a program that would be able to track the moving visual stimuli (white dots on a dark background) while also simultaneously track moving animals (dark tadpoles on an alternating white/dark background), and then overlay the two and report both encounters and avoidance events. While it is possible to filter out the visual stimuli so that the tadpoles can see them but the camera does not, doing so would preclude the investigator from being able to manually check the data and it would exclude any post-hoc re-analysis of years of pre-existing data. To address these needs, we developed X-Tracker, a Matlab code that digitally extracts the visual stimuli from the video data, tracks the tadpoles, identifies the stimuli, and then subsequently overlays the visual stimuli back onto the tadpole tracks to score encounters and avoidance events.

As shown in the flowchart (Figure 3), X-Tracker first calculates the mean background and then subtracts the stimulus dots from each frame. To obtain the mean background, the program averages the dot-containing frames together, and then subtracts each frame from this mean background so that only moving objects (dots and tadpoles) remain. It removes the stimulus dots by thresholding, as the tadpoles are darker than the mean background while the dots are lighter. The program then uses Blob Filtering methods contained within Matlab to extract the locations of the tadpoles based on the density of their gut and assigns each tadpole a location based on distance to the next location. At each frame, for each extracted tadpole, X-Tracker uses Kalman filtering to predict where the tadpole may swim in the subsequent frame. It then uses the Hungarian algorithm to connect tadpole detections in this subsequent frame to tadpoles in the current frame based on these predictions, thus creating a continuous swimming trajectory for each tadpole. Animals are only scored if they are moving at a minimum rate of 50 pixels/second. Tadpoles must be moving to be scored for an avoidance response. Any animal that is not moving will be considered part of the background and will not be assigned or scored.

**Figure 3.**
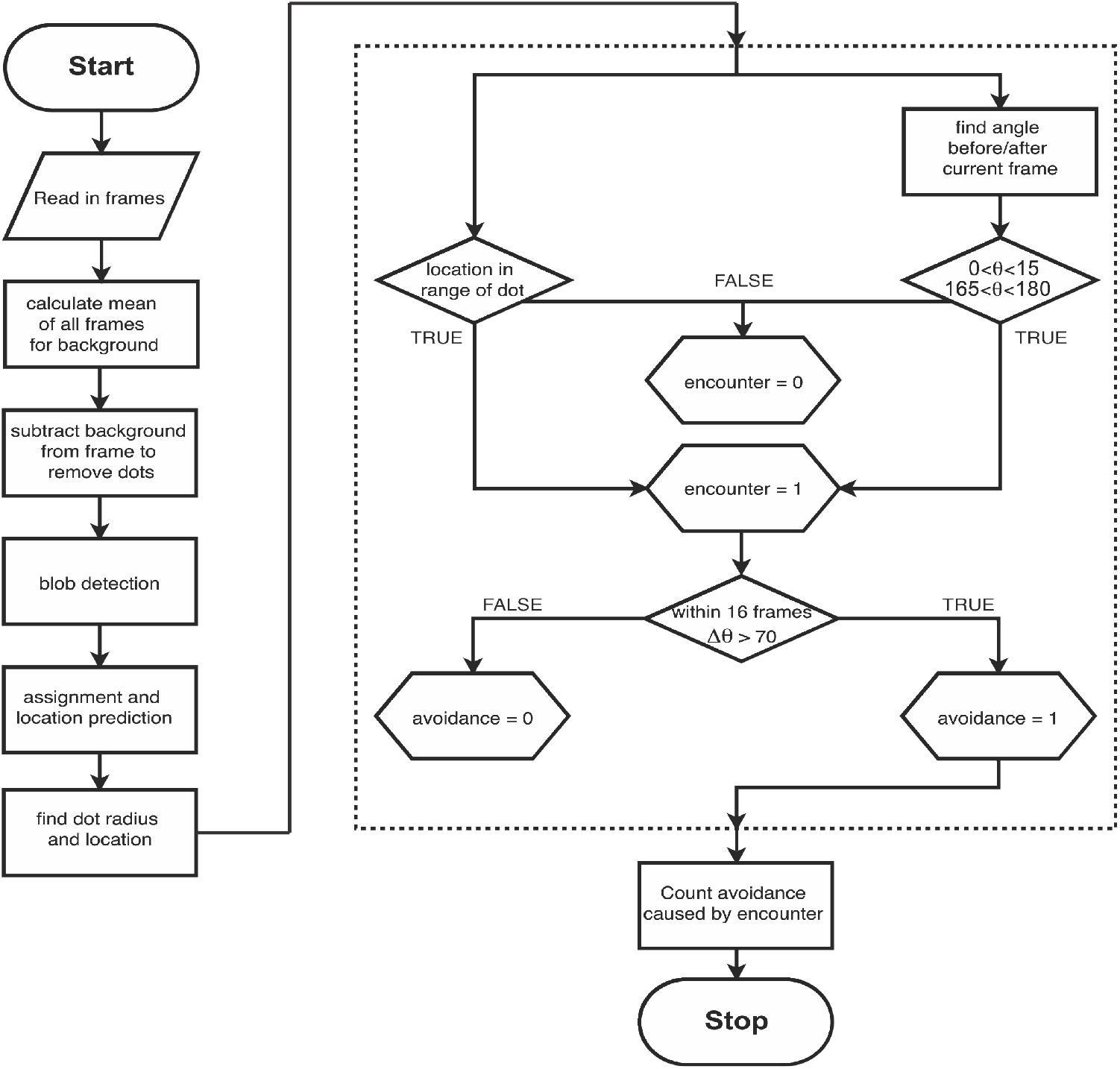
Computational flow-chart of X-Tracker automated behavior analysis code. The left side of the flow-chart describes the preliminary sequence of steps X-Tracker performs to obtain the coordinates of the dots and tadpoles at each video frame. The boxed portion depicts the logic used to assess when an encounter occurs and how to score it when one does occur. A tadpole must both be within range of a dot and be swimming at an angle within 15° degrees of perpendicular to the dot’s trajectory in order for an encounter to be scored; the tadpole must then turn 90±20° from its prior trajectory for the encounter to be scored as an avoidance. Total avoidances and encounters are summed at the end of the program.

X-Tracker determines the location of radius of each oncoming stimulus dot using the original video. Images annotated with the output of these calculations are shown as a biological flowchart in Figure 4. Stimulus dots are presented in a random array at a uniform size and speed, and are detected by X-Tracker via image thresholding to extract the lighter dots followed by morphological opening with a disk-shaped structuring element. The dot presentation software is also available in the Cline Lab GitHub repository. Partial dot artifacts created by edge-effect diffraction are filtered out using a circularity threshold so that only round stimuli are tracked (Figure 4, asterisks). Using the locations of the tadpoles and the locations and radii of the dots, the data is analyzed to identify frames in which a tadpole comes into contact with, or encounters, a dot.

**Figure 4.**
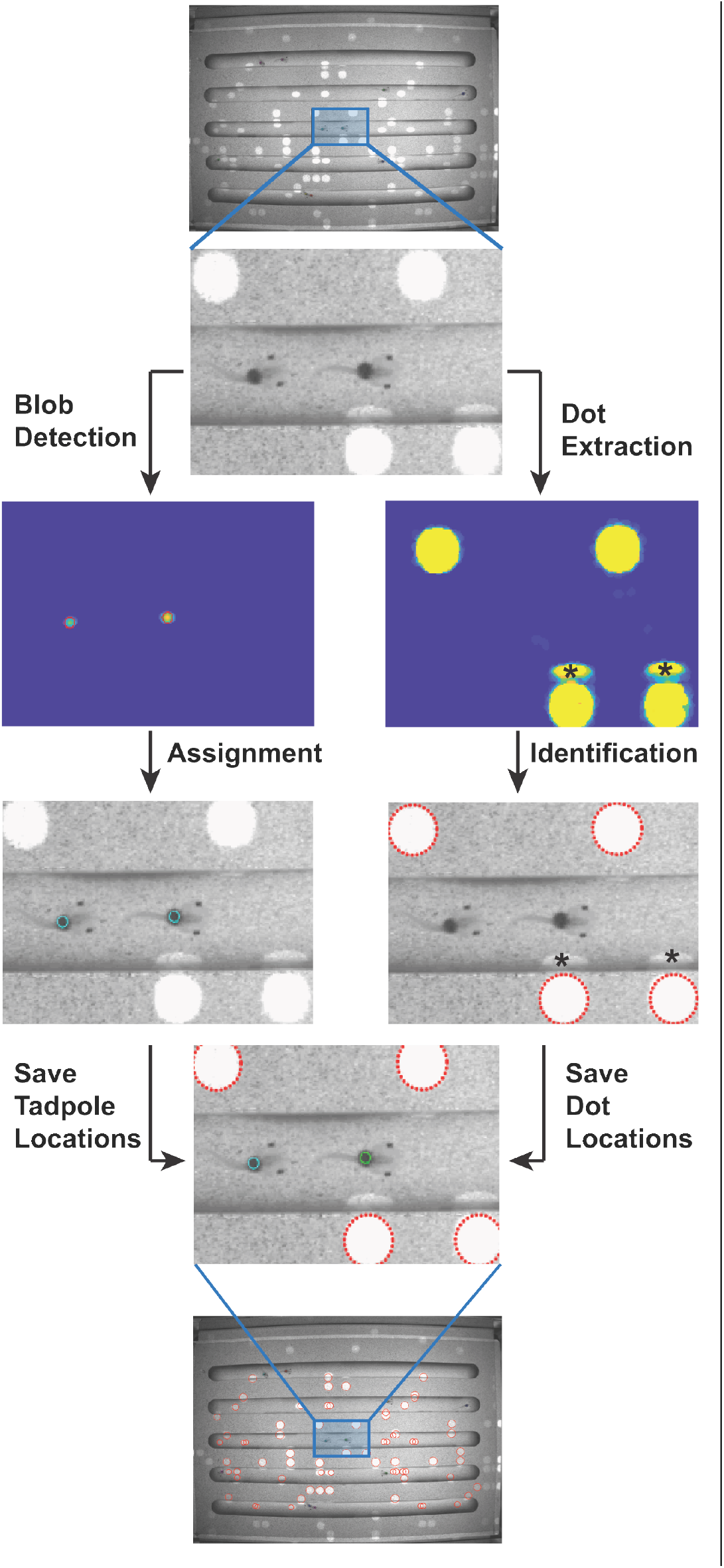
Biological flow-chart of X-Tracker automated analysis. Images are shown from different stages throughout the automated analysis. A small field of view is shown for presentation purposes. The dots are extracted, and tadpoles are identified by Blob Detection methods. Locations for both dots and tadpoles are identified, saved, and then overlaid to determine the number of encounters and the fraction of avoidance events. Asterisks denote partial dot artifacts created by edge-effect diffraction that are subsequently filtered out using a circularity threshold in the software, as indicated by the dotted red outlines.

Encounters between a tadpole and a dot are determined as follows. To estimate the co-ordinates of the field of view, X-Tracker expands the previously determined tadpole location to include the eyes. We determined that at stage 47, the average distance between the tadpole’s gut and eyes is approximately 15 pixels, or ∼250um. Using this distance as a radius, points in a semicircle around the head and eyes are defined and assigned as new coordinates for the tadpoles. This value can be adjusted to accommodate tracking of larger or smaller animals as needed. At stage 47 (Nieuwkoop and Faber, 1967), when tadpoles first begin to exhibit a visual avoidance response (Dong et al., 2009; McKeown et al., 2013), the visual focal distance is approximately 250um (Richards et al., 2012), therefore we set the software to define an encounter when a dot and an eye are juxtaposed, in adjacent pixels.

Once the program has detected a tadpole-dot encounter, it will then determine whether the animal exhibited an avoidance reaction in response to the dot stimulus. Based on set criteria from previous manual analysis protocols, we define an avoidance as a change in trajectory within 500ms of an encounter between the eye of a moving tadpole and a perpendicularly approaching dot (Dong et al., 2009; McKeown et al., 2013; Shen et al., 2011). In both the open field and the channel chambers, an avoidance response is scored when it results in a turn of 90±20 degrees (Figure 2A, B) (Dong et al., 2009; McKeown et al., 2013; Shen et al., 2011).

To facilitate the usage of X-Tracker for high-throughput analysis, we have streamlined the analysis of multiple videos with a batch input function. Utilizing this function, X-Tracker can be run within a folder containing multiple video recordings. The program will then execute with same preset settings for each file in sequence, saving the results in a single summary file. This batch input feature allows for the automated analysis of a large number of videos without any additional input required from the user for each file. The batch input code for X-Tracker is also available in the Cline Lab GitHub repository.

To validate the data generated by X-Tracker, we tested a manually scored dataset against the code. The avoidance index for this clutch was identical between the human manual scoring with an average ± SEM of 0.214±0.024 for manual vs. 0.202±0.036 for X-

Tracker (Figure 5). The difference lies in the number of events that can be scored using the X-Tracker automated analysis (Table 1). As in previous experiments, the human scoring was limited to the first 10 encounters (Gambrill et al., 2019; McKeown et al., 2013; McKeown et al., 2017; Shen et al., 2011), scoring between 6-10 events per animal in this particular dataset, whereas X-Tracker was unlimited, counting from 6 to 34 events per animal (Figure 5 and Table 1). It is important to note X-Tracker has more stringent movement criteria, requiring animals spend less than 50% of the video in a single spot, therefore more animals are excluded from the analysis than by manual analysis (Figure 5). However, this creates a consistent and reproducible analysis that is not subject to user bias. Nonetheless, X-Tracker returns avoidance behavior data that is consistent with manual analysis.

**Figure 5.**
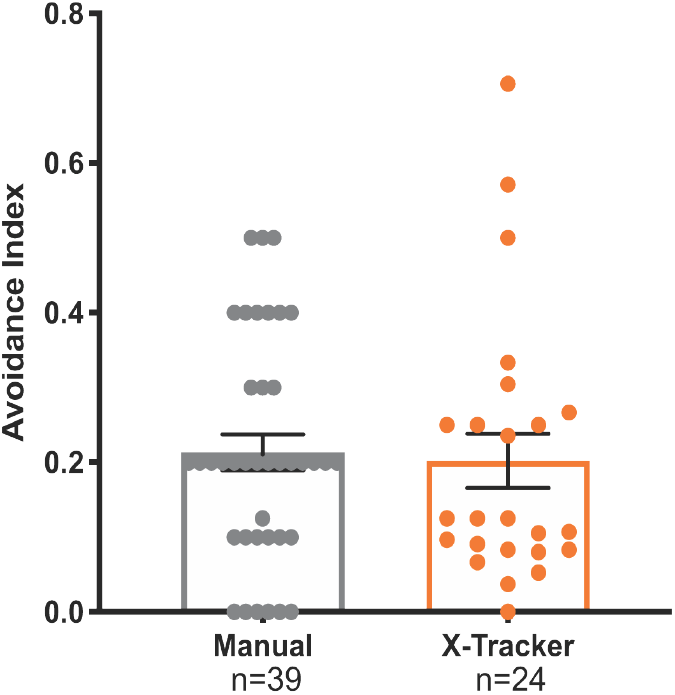
X-Tracker automated analysis is similar to manual human analysis. Avoidance indexes comparing Human Manual Analysis (gray) vs. X-Tracker Automated Analysis (orange) on identical videos. Data are shown as average ± SEM overlaid with individual data points. There is no significant difference in the averages between the groups by ANOVA with post hoc Tukey’s for multiple comparisons.

## Discussion

Here we present X-Tracker, an automated behavior tracking code that can detect both moving visual stimuli and free-swimming animals, score encounters between animals and stimuli, and report avoidance events. X-Tracker reproducibly standardizes behavior analysis, generating results indistinguishable from manual analysis. X-Tracker is hands-off, high-throughput, and makes behavior analysis objective and unbiased. To increase the number of animal encounters with the visual stimuli, we created a new channel chamber for testing behavior. This channel system forces animals to swim perpendicularly to the moving stimulus, thereby increasing the frequency of visual encounters. And lastly, we improved behavior rig illumination with LED strips and upgraded the detection camera, both of which generate higher quality uniform images to optimize input for automated detection software. Together these advancements increase the detection of visual encounters and avoidance events, decrease potential inter-investigator variance in scoring events, and significantly decrease time-consuming human involvement in the analysis, making visual avoidance behavior assays more accessible for researchers.

Visually guided behaviors have been used to investigate brain function in various systems, yet automated analysis of certain visually guided behaviors has been difficult. While animal tracking analysis is relatively straight-forward (Gal et al., 2020; Kohlhoff et al., 2011), and analysis packages are commercially available (Noldus EthoVision and Viewpoint ZebraLab being the most commonly used), simultaneous tracking of both animals and visual stimuli in naturalistic conditions has proven more difficult (Michaiel et al., 2020). In *Xenopus*, researchers have developed creative low through-put ways around this hurdle by testing one animal at a time (Khakhalin, 2020; Khakhalin et al., 2014). In other free-swimming species such as the Zebrafish, investigators have used filters to block out visual stimuli so that they only track the animals’ movements in response to a non-random stimulus, making the automated analysis simpler, but limiting the variety of stimuli that can be used and creating additional concerns about habituation to the stimulus (Larsch and Pantoja, 2019; Lowe, 1987; Randlett et al., 2019). X-Tracker allows a variety of stimuli to be tested (size, speed, shapes, patterns) with ease. In addition to visual avoidance behavior, *Xenopus* tadpoles exhibit schooling behavior (Truszkowski et al., 2016), wall-following behavior (Hanzi and Straka, 2018), startle responses (James et al., 2015), color preference (Moriya et al., 1996), and seizures (James et al., 2015). X-Tracker can be easily adjusted to automate the analysis of this increasing repertoire of *Xenopus* behaviors, aiding in the investigation of brain function in a developing system. Lastly, X-Tracker can be readily modified to detect larger (and smaller) animals, making this software applicable to not only different developmental stages of *Xenopus laevis*, but also other species including *Xenopus tropicalis* and other amphibian tadpoles, Zebrafish, and teleost fishes. Because the background is subtracted, X-Tracker can detect animals in any transparent chamber, making it adaptable to endless sizes of animals and visual stimuli.

X-Tracker allows for unbiased, hands-free automated analysis of *Xenopus* visual avoidance behavior. Previously, the rate-limiting factor for these experiments was the time-consuming manual analysis. However, X-Tracker frees up the analysis time, allowing for large high through-put experiments testing molecular pathways via drugs and genetic contributions to behavior. We expect that this software will significantly aid researchers in designing experiments to identify the molecular and cellular components related to brain development and behavior in *Xenopus* and other free-swimming animals.

### Information Sharing Statement

Software presented here is publicly available for non-commercial use in the Cline Lab GitHub repository at https://github.com/Cline-Lab/Tadpole-Behavior-Automation. Note: Early versions of the X-Tracker software code were originally called TAD9000 which may be the title of some older drafts on GitHub. 2.

The datasets generated during and/or analyzed in the current study are available from the corresponding author on reasonable request.

## Acknowledgements

We thank members of the Cline lab, past and present, for helpful discussions, Lin-Chien Huang for making the first channel prototype, Steve Barry at the Salk Institute instrument lab for making the channel chambers, and all the students, including but not limited to Tyler Wishard, Heidi Sharipova, Breanna Gomez, and Ila Peeler, who painstakingly scored behavior manually prior to this development. This work was supported the US National Institutes of Health (EY011261 and EY027437 to HTC, and P30 EY019005 to J. Reynolds at the Salk Institute), and an endowment from the Hahn Family Foundation to HTC.

## Authors’ contributions

Conceptualization (HTC, CRM, ACT, CLM), Formal Analysis (CRM, ACT), Funding Acquisition (HTC), Investigation (NM, KA), Methodology (CLM), Software (CLM, ACT), Supervision (HTC), Visualization (CRM), Writing – original draft (CRM), Writing – reviewing & editing (HTC, CRM, ACT, NJM).

## Conflicts of interest/Competing interests

The authors have no conflicts of interest or competing interests to disclose.

## Notes

### Competing Interest Statement

The authors have declared no competing interest.

